# Somatic gonad morphogenesis in *C. elegans* requires the heterochronic pathway acting through HBL-1

**DOI:** 10.1101/2024.11.11.623063

**Authors:** Madeleine Minutillo, Kevin Kemper, Maria Ivanova, Erika Pianin, Eric G. Moss

## Abstract

Characteristically, the heterochronic genes control the stage-specific timing of developmental events throughout the developing *Caenorhaditis elegans* larva while gonad development occurs mostly normally. In a few cases, some timing regulators lead to detectable variations in gonad development with little to no effect on fertility. We found that a double mutant of a *lin-28* null allele and *hbl-1* hypomorphic allele results in a catastrophic failure of gonad morphogenesis resulting in sterility. This defect includes a high-penetrance disruption of normal gonad arm migration as well as frequent absence of one or both spermathecae. We demonstrate that the abnormal gonad morphology and novel sterility phenotype is ultimately due to loss of *hbl-1* activity in larval development. The mechanism of how *lin-28* positively influences *hbl-1* activity is unresolved. We demonstrate a direct interaction between the RNA-binding protein LIN- 28 and the 5’UTR of *lin-46*, and in turn, a direct interaction between LIN-46 and two zinc fingers of HBL-1. Nevertheless, our genetic analysis indicates that *lin-46* accounts for only part of *lin-28*’s regulation of *hbl-1* and that some portion of *lin-28*’s effect is independent of *lin-46*.

## Introduction

The heterochronic genes of the nematode *Caenorhabditis elegans* comprise a well-characterized regulatory system explicitly controlling timing during tissue and organ formation in animals (Moss 2007; Rougvie and Moss 2013). This genetic pathway governs the succession of cell division and differentiation events in several tissues during *C. elegans* larval development. Most tissues in this animal that develop post-embryonically execute specific developmental events at each larval stage. Mutations in heterochronic genes cause stage-specific events to be either skipped or repeated, resulting in precocious or reiterative phenotypes, respectively. The most significant consequence of abnormal timing for the worm is an asynchrony between the development of the egg laying system, which emerges from the hypodermis and other tissues, and the gonad development, which occurs on its normal schedule in these mutants.

The majority of heterochronic mutants are fertile with properly formed and fully functional gonads. Certain mutations cause notable defects in the path of migration of the distal tip cells (DTCs) as the gonad grows and takes shape (Tennessen et al. 2006; Cecchetelli and Cram 2017). For example, particular alleles of *daf-12* cause both reiterative hypodermal development as well as an abnormal gonad shape where the DTCs take aberrant paths during gonad growth (Antebi et al. 1998; Hammell et al. 2009). The DTCs normally turn dorsally at the third larval stage (L3) molt but often fail to do so in these mutants. Despite the deviant migration, *daf-12* mutant animals have undisturbed fertility. *daf-12* encodes a nuclear hormone receptor that regulates the transcription of microRNA genes involved in heterochronic gene regulation (Hammell et al. 2009).

Choi and Ambros documented LIN-28’s role in the normal morphogenesis of the hermaphrodite spermatheca, the part of the somatic gonad where sperm are held and fertilization takes place (Choi and Ambros 2019). Mutations in genes causing structural abnormalities that affect ovulation, fertilization, spermathecal exit and egg laying reduce the overall fertility of an animal (Kovacevic and Cram 2010; Choi and Ambros 2019).

Choi and Ambros also observed that RNAi of *hbl-1* caused a similar spermathecal defect. It is not clear however how *lin-28* or *hbl-1* affect spermathecal development or whether that effect is related to their roles in the developmental timing pathway.

Mutations in either *lin-28* or *hbl-1* causes widespread precocious hypodermal development, where events of the second larval stage (L2) are skipped (Ambros and Horvitz 1984; Abrahante et al. 2003; Lin et al. 2003). LIN-28 is a conserved RNA-binding protein that controls events of the L2 stage via post-transcriptional regulation of *hbl-1* at least in part via the negative regulation of *lin-46* (Pepper et al. 2004; Vadla et al. 2012; Ilbay and Ambros 2019). A second activity of LIN-28 blocks the maturation of the pre-let-7 microRNA by which it controls events of the L3 stage (Vadla et al. 2012). HBL- 1 is a zinc finger transcription factor belonging to the Ikaros family (Fay et al. 1999). Null alleles of *hbl-1* die as late-stage embryos or young larvae because *hbl-1* is required for hypodermal differentiation late in embryogenesis (Fay et al. 1999; Abrahante et al. 2003; Lin et al. 2003). Hypomorphic alleles of *hbl-1* have been found that survive to adulthood but show precocious heterochronic defects, specifically skipping events of the L2, much like *lin-28* null alleles (Abrahante et al. 2003; Lin et al. 2003). *hbl-1* is negatively regulated by let-7 family microRNAs acting through its 3’UTR (Abbott et al. 2005).

Numerous genetic studies have placed *hbl-1* downstream of *lin-28*, and it is likely the most direct regulator of L2 fates of the known heterochronic genes. Because null alleles of *hbl-1* die as embryos, studying its postembryonic role has been a challenge that has relied on hypomorphic alleles, therefore, some genetic tests that typically rely on null alleles have been difficult to evaluate. *hbl-1* activity in the heterochronic pathway appears to be regulated both post-transcriptionally and post-translationally. Through its 3’UTR, *hbl-1* expression is regulated by members of the let-7 family of microRNAs, where deletion of either the microRNAs or the 3’UTR causes misexpression of the protein. Furthermore, LIN-46 appears to affect HBL-1 localization (Vadla et al. 2012; Ilbay and Ambros 2019). Here we provide evidence for a direct association between LIN-28 and the *lin-46* mRNA as well as between LIN-46 and HBL-1 that appears to alter HBL-1’s activity.

Because *lin-28* and *hbl-1* both control L2 events, we attempted to sort out whether *lin-28* and *hbl-1* act in a linear pathway to control hypodermal development or whether the pathway branches at some point. We made an unexpected observation concerning gonad morphogenesis that nevertheless helps us answer that question.

## Results

### Fertility requires the combined activities of lin-28 and hbl-1

The genes *lin-28* and *hbl-1* are required for normal L2 development, with mutations in each causing animals to skip L2-specific events in the hypodermis (Moss et al. 1997; Abrahante et al. 2003; Abbott et al. 2005). As part of addressing the relationship between these two regulators, we attempted to construct a double mutant between a null allele of *lin-28* (*ga54*) and a hypomorphic allele of *hbl-1* (ve18) that shows only a heterochronic phenotype (Abrahante et al. 2003). We found that animals homozygous for both alleles were sterile and lacked the characteristic protruding vulva of each alone (Fig. 1). Occasionally, some double mutants were able to produce offspring, but the fertility and fecundity of these animals was severely reduced relative to the single mutants alone (Table 1, lines 1-6, columns A and B). However, we could maintain a balanced strain containing both alleles when *lin-28(ga54)* was homozygous and *hbl-1(ve18)* was heterozygous; all double mutants we characterized were offspring of such balanced strains.

**Fig. 1.**
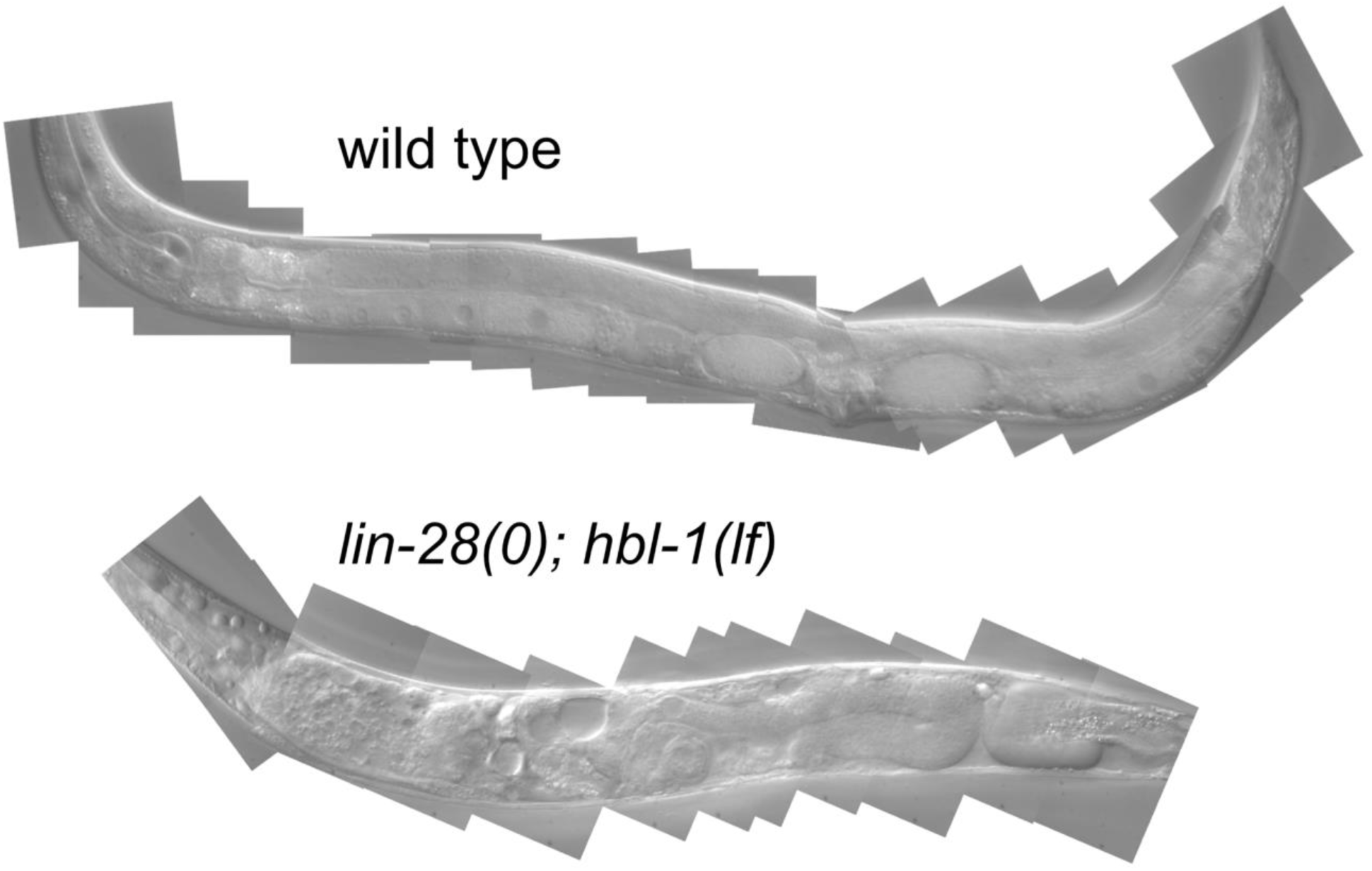
**Gross terminal phenotype of *lin-28; hbl-1* double mutant of *C. elegans.* Top**, Wildtype *C. elegans* adult with mature oocytes and two eggs in the uterus. Anterior to the left and dorsal up. **Bottom**, a similarly-aged *lin-28(0); hbl-1(lf)* animal, displaying an overall “dumpy” morphology, apparent lack of vulva, disorganized gonad, vacuoles, yolk deposits, and sterility.

**Table 1.** Phenotypes of mutant strains

|  | Genotype <sup>a</sup> | A<br>% fertile | B<br>Average brood size of fertile Animals (n) <sup>b,d</sup> | C<br>Number of seam cells at adulthood (n) <sup>b</sup> |
| --- | --- | --- | --- | --- |
| 1 | wild type | 100 | 262.7 (10) <sup>+</sup> | 16.1 (30) |
| 2 | *lin-28(ga54)* | 100 | 20.3 (57) | 10.8 (28) |
| 3 | *hbl-1(ve18)* | 95 | 48.4 (44) | 20.9 (27) <sup>c</sup> |
| 4 | *hbl-1(ae23)* | 100 | 248.9 (24) <sup>+</sup> | 16.1 (31) |
| 5 | *lin-28(ga54); hbl-1(ve18)* | 10.6 | 3.4 (66) | 10.4 (21) |
| 6 | *lin-28(ga54); hbl-1(ae23)* | 19 | 1.92 (68) | 11.1(22) |
| 7 | *lin-28(ga54); hbl-1(ae26)* | 100 | 24.4 (70) | 11.1(41) |
| 8 | *lin-28(ga54); lin-46(ma174); hbl-1(ve18)* | 65 | 4.5 (40) | 12.6 (46) |
| 9 | *lin-28(ga54); lin-46(ma174); hbl-1(ae23)* | 92 | 10.7 (62) | 15.8 (37) |
| 10 | *lin-46(ae30)* | 100 | 56.5 (36) | 12.1 (31) |
| 11 | *lin-46(ae30); hbl-1(ve18)* | 82.8 | 9.6 (35) | 13.5 (35) |
| 12 | *lin-46(ae30); hbl-1(ae23)* | 100 | 48.4 (35) | 12.2 (34) |
| 13 | *hbl-1(AID)*, no auxin | 100 | 323.5 (10) <sup>+</sup> | 16.1 (36) |
| 14 | *hbl-1(AID)* | 100 | 51.1 (15) | 14.1 (36) |
| 15 | *hbl-1(AID+RNAi)*, no auxin | 91 | 12.1 (43) | 12.2 (24) |
| 16 | *hbl-1(AID+RNAi)* | 50 | 6.9 (36) | 11.8 (26) |
| 17 | *lin-28(ga54); hbl-1(AID+RNAi)* | 0 | 0 (39) | 10.9 (39) |
| 18 | *lin-28(ga54); lin-46(ma174); hbl-1(AID+RNAi)* | 27 | 2.9 (30) | 13.5 (35) |

<sup>a</sup> All strains examined were homozygous for alleles indicated and carry an integrated transgene
*wls78(ajm-1::GFP +SCMp::GFP)* to mark seam cells.
<sup>b</sup> Number of animals (n) scored for each genotype is indicated in parenthesis.
<sup>c</sup> *hbl-1(ve18)* animals have a reduced number of seam cells from the L2 onward compared to wild type, indicative of skipping L2-specific cell divisions, but have been previously reported to produce some extra seam cell nuclei at the end of larval development (Abrahante et al. 2003; Abbott et al. 2005). This is the reason for the high seam cell count for adults of this strain.
<sup>d</sup> All strains examined were egg laying defective unless otherwise labeled with “+” superscript denoting wild type egg laying capability.

Like the single mutants, the *lin-28; hbl-1* animals showed a precocious heterochronic phenotype characterized by a reduced number of seam cells and formation of adult alae at the end of the L3 (Table 1, lines 1-6, column C). The number of seam cells in the double mutant was not appreciably reduced compared to either single mutant. Similarly, the double mutant displayed precocious alae like that of each single mutant: patches of alae appearing at the L2 molt and full-length adult alae at the L3 molt (<u>Fig. S1</u>). However, unlike single mutants the double mutant displayed molting defects, becoming trapped in the unshed cuticle. Occasionally, when a few progeny were produced they can be observed trapped in the deceased hermaphrodite’s cuticle (<u>Fig. S2</u>). As in *lin-28* mutants, vulval development in *lin-28(0); hbl-1(lf)* animals began one stage early, VPCs dividing at the L2 molt. But unlike *lin-28* mutants, vulva development did not complete and appeared to arrest during morphogenesis. This explains the lack of protruding vulva seen in the single mutants where the vulva completes morphogenesis one stage early (Euling and Ambros 1996; Abrahante et al. 2003) (<u>Fig. S1</u>). Arrested vulva development has been observed in animals grown on *hbl-1* RNAi (Fay et al. 1999). Also like the *lin-28* single mutant, formation of one or more pseudovulva was typical, although they too arrested mid morphogenesis (<u>Fig. S1</u>).

To test whether a milder version of this phenotype might result from using a weaker allele of *hbl-1*, we characterized the phenotype of a double mutant containing *hbl-1(ae23).* The *hbl-1(ae23)* allele has a small deletion at the 3’ end of the open reading frame that disrupts the two C-terminal dimerization zinc fingers of the protein. The *hbl-1(ae23)* allele produces no apparent phenotype on its own (Table 1, line 4). Surprisingly, the *lin-28(ga54); hbl-1(ae23)* double mutant displayed a strong sterility phenotype, albeit slightly less penetrant than that of *lin-28(ga54); hbl-1(ve18)* (Table 1, line 6). For all other traits, including heterochronic phenotype as well as gross body morphology, the *lin-28(ga54); hbl-1(ae23)* double mutant resembled the *lin-28(ga54); hbl-1(ve18)* double mutant.

### LIN-28 and HBL-1 are needed for gonad morphogenesis

To explore the basis of this sterility, we examined gonad development in *lin- 28(0); hbl-1(lf)* double mutants. The gonad appears normal through the L2 when the first heterochronic defects of these genes appear. Beginning in the L3, defects in gonad development become apparent: the gonad arms do not follow their normal reflexive path to the center of the animal. By the late L4 the malformation of the uterus is observable. By adulthood, the gonads of these animals appeared disorganized with significant accumulation of yolk in the pseudocoelomic space. This pooling of yolk within the pseudocoelom indicates additional defects in the double mutant animals. It is possible that there are further abnormalities of the somatic gonad in which sheath cells surrounding the proximal gonad are unable to transport yolk from the intestine to the oocytes.

To quantify gonad migration, we used a *lag-2::GFP* reporter which marks the distal tip cells (DTCs) as the gonad grows and migrates during the larval stages (Blelloch and Kimble 1999). Through the L2, DTC shape, position, and movement appeared normal in *lin-28(0); hbl-1(lf)* animals. Beginning in the L3, when the DTCs normally turn dorsally and reverse their direction along the anterior-posterior axis, the DTCs in *lin-28(0); hbl-1(lf)* animals migrated abnormally, often continuing toward the head or tail, and this wayward migration continues into the L4 (Fig. 2). This deviation in migration was the first visible defect identifiable in this double mutant’s gonad development. Abnormal migration of the DTC of one arm was displayed by 24% of animals, abnormal migration of the DTCs in both arms was displayed by 54% of animals (Table 2).

**Fig. 2.**
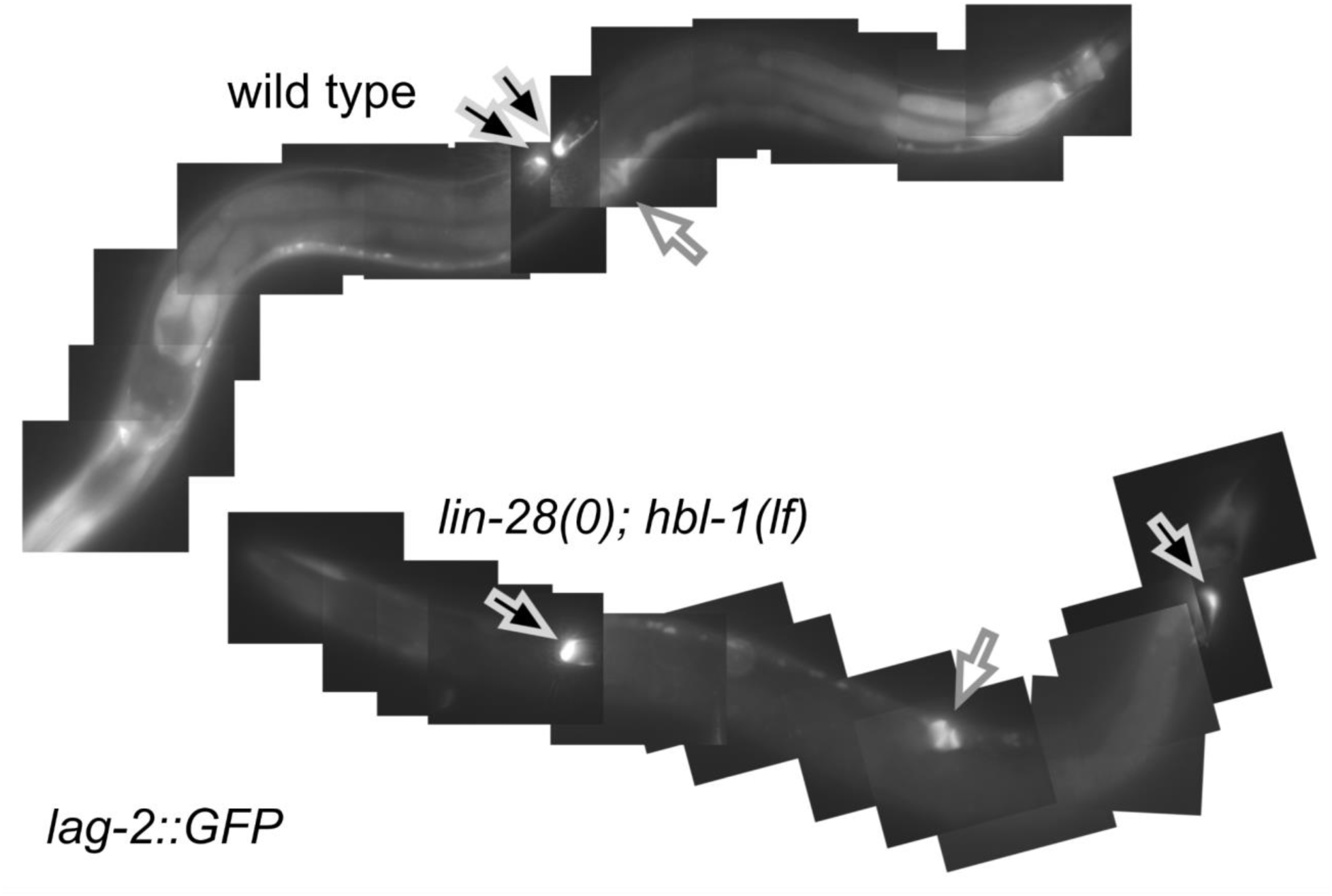
Abnormal DTC migration in *lin-28; hbl-1.* Top, Wildtype *C. elegans* adult expressing *lag-2::GFP.* Black arrows, locations of fluorescing distal tip cells (DTCs). White arrows, vulva. The DTCs are located near one another on the dorsal side opposite the vulva. **Bottom,** a similarly-aged *lin-28; hbl-1* double mutant, also expressing *lag-2::GFP*. The DTCs are located far from each other near the head and tail.

**Table 2.**
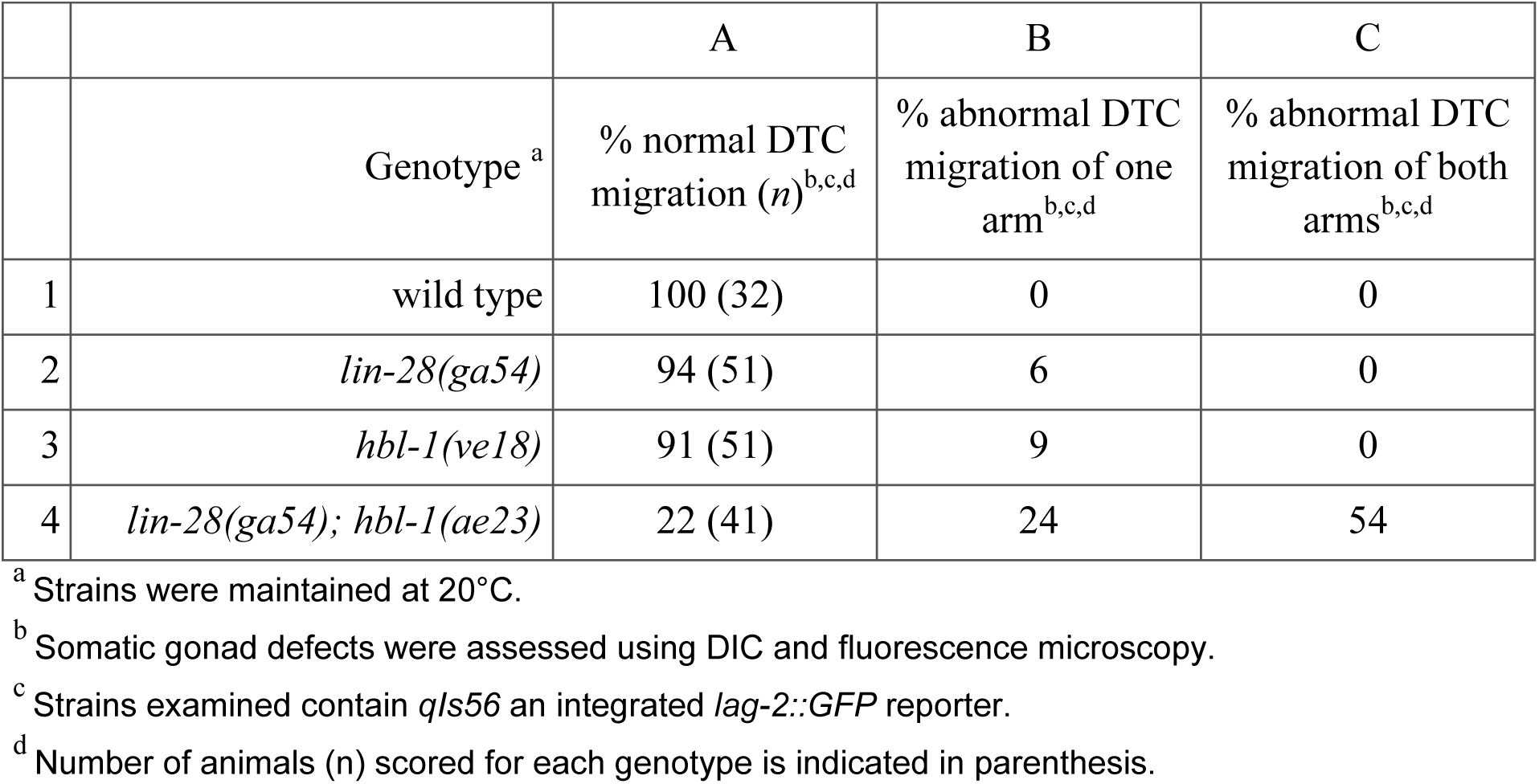
Abnormal DTC migration in single and double mutants

The sterility phenotype observed in the double mutants could be due to defects in spermathecal development (Choi and Ambros 2019). We observed that although oocytes are formed in these double mutant animals, few gained entry into the spermatheca. The oocytes that were able to enter the spermatheca were unable to exit the spermatheca. To assess spermathecal development, we examined *lin-28(0); hbl- 1(lf)* double mutants using a *fkh-6::GFP* reporter (Chang et al. 2004). This reporter is expressed in the spermatheca as well as spermathecal precursor cells. In wildtype animals, the reporter is consistently expressed in both spermathecae and displays the typical tube shape assumed by the spermatheca at the end of development (Fig. 3)(Chang et al. 2004).

**Fig. 3.**
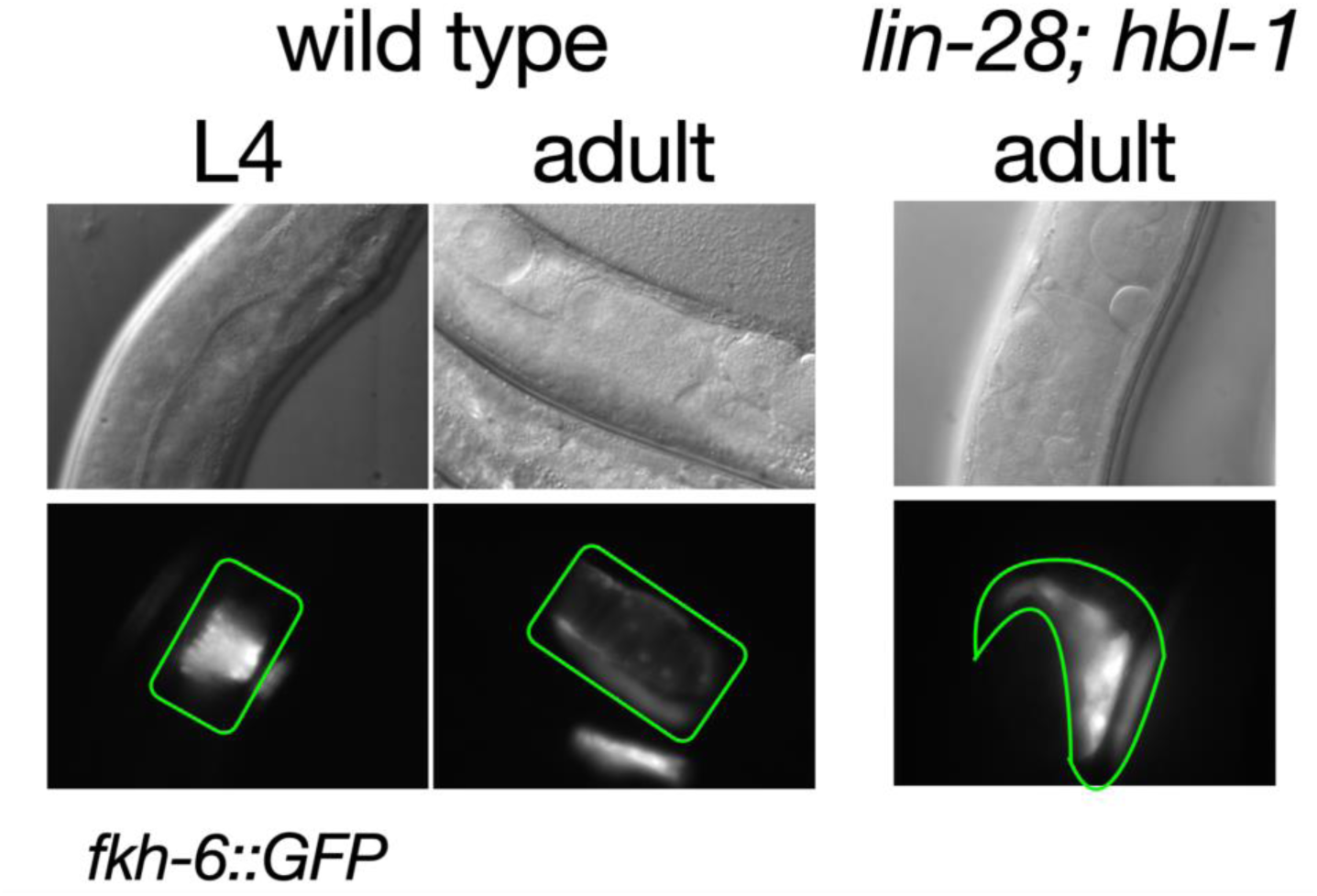
Abnormal spermatheca formation in *lin-28; hbl-1.* Left, Wild type *C. elegans* L4 and adult expressing *fkh-6::GFP* in the developing and mature spermathecae, respectively. **Right,** adult *lin-28; hbl-1* double mutant showing an incompletely developed and abnormally shaped spermatheca. Black arrow indicating mass of oocytes in proximal gonad.

In the *lin-28(0); hbl-1(lf)* double mutant, the presentation of the reporter was markedly different from wild type in two ways. First, the fluorescent signal was variable, present in either both, one, or neither arms of the gonad (Table 3). As best that could be determined, the variability in reporter expression was not due to reporter failure. The absence of reporter expression was never seen in either single mutant alone (Table 3). In the double mutant, gonad arms that displayed little to no reporter expression were also observed to be abnormal in appearance. The area of the gonad arm at which the spermatheca should be present appeared to resemble that of an earlier stage of development, in which conspicuous features of the spermatheca were not observable. When the spermatheca did not form, sometimes only a few (3-12) of the 24 cells that make up the spermatheca could be seen expressing the reporter. This suggests failure of spermathecal development at a stage prior to morphogenesis.

**Table 3.** *fkh-6::GFP* reporter expression in single and double mutants

|  |  | A | B | C |
| --- | --- | --- | --- | --- |
|  | Genotype <sup>a</sup> | % with 2 fluorescent spermathecae (*n*) <sup>b,c,d</sup> | % with 1 fluorescent spermatheca <sup>b,c,d</sup> | % with 0 spermathecae <sup>b,c,d</sup> |
| 1 | wild type | 100 (63) | 0 | 0 |
| 2 | *lin-28(ga54)* | 100 (62) | 0 | 0 |
| 3 | *hbl-1(ve18)* | 100 (62) | 0 | 0 |
| 4 | *lin-28(ga54); hbl-1(ae23)* | 44 (62) | 38 | 18 |

<sup>a</sup> Strains were maintained at 20°C.
<sup>b</sup> Somatic gonad defects were assessed using DIC and fluorescence microscopy.
<sup>c</sup> Strains examined contain *ezls2* an integrated *fkh-6::GFP* reporter.
<sup>d</sup> Number of animals (*n*) scored for each genotype is indicated in parenthesis.

The second way the reporter appeared different in the mutant is that reporter- expressing spermatheca had an irregular morphology instead of the wild type tube shape (Fig. 3). It sometimes exhibited what looked to be a cross-section of the spermatheca rather than the left or right side, possibly indicating improper orientation of the tissue. When the spermatheca did form correctly, sperm could be observed within it by adulthood. In most cases, the proximal gonad of adult mutant became clogged with a mass of oocytes and oocyte fragments (Fig. 3). This observation indicated the primary cause for the failure of fertilization of mature oocytes was defective spermathecal development. In addition to spermathecal defects, *lin-28(0); hbl-1(lf)* double mutants also displayed malformed uteri. Analysis of vulva morphology by DIC microscopy revealed that morphogenesis ceased after vulval cell invagination but before anchor cell invasion. Consequently, the formation of the uterine seam cell (utse) did not occur (Gupta et al. 2012).

Because *lin-28* and *hbl-1* act during early stages of larval development, we wanted to examine the effect their loss would have on the early stages of somatic gonad development. We used a strain containing the *ckb-3p::mCherry::H2B* reporter, which is expressed in the daughters of the gonad precursor cells Z1 and Z4 in the first and second larval stage (Tenen and Greenwald 2019). We compared the number of fluorescent cells in the L1 and L2 stages of wildtype animals to *lin-28(0); hbl-1(lf)* double mutant and found no difference. While we were not able to see a cell lineage defect in these particular cells of the somatic gonad, it is possible the lineage defect occurs in other cells formed in the early larval stages or the defect simply occurs at a later developmental stage. It is also possible the defect affects the identity or behavior of certain cells. This would not affect cell number or location initially but morphological defects would result later in development.

Given that the loss of *lin-28* and *hbl-1* together dramatically impacts hermaphrodite somatic gonad development, we reasoned the male somatic gonad would also be affected. To address this question, we generated and examined males of the *lin-28(0); hbl-1(lf)* double mutant strain containing the *lag-2::GFP* reporter, which is expressed in the linker cell that leads gonad migration in males. While the loss of *lin-28* and *hbl-1* activity causes deformities in the male tail due to heterochronic defects, surprisingly, we observed no abnormal gonad migration in males.

### LIN-28 binds directly to lin-46’s 5’UTR

To gain a mechanistic understanding of the relationship between *lin-28* and *hbl- 1*, we further characterized the role of the protein LIN-46 in the heterochronic pathway (Pepper et al. 2004). Genetic evidence suggests that *lin-46* acts downstream of *lin-28* and upstream of *hbl-1* (Abbott et al. 2005). Mutations in *lin-28* and *lin-46* suppress each other, suggesting either that *lin-28* acts partly via *lin-46* or that they act in parallel (Pepper et al. 2004). A recent study showed that mutations in *lin-46’s* 5’UTR cause a phenotype similar to *lin-28* loss-of-function, showing that the RNA-binding protein LIN- 28 acts at least in part by directly regulating *lin-46* expression (Ilbay et al. 2021).

We tested whether LIN-28 can directly interact with the *lin-46* mRNA by performing the yeast three-hybrid assay, as we had done previously to show LIN-28 binding to pre-let-7 and other pre-microRNAs (Vadla et al. 2012). We found that LIN-28 could bind the 35-nucleotide wildtype *lin-46* 5’UTR sequence, but not to a mutant 5’UTR containing a 6-nt deletion corresponding to allele *ae30* (Table 4). *lin-46(ae30)* is one of several mutations in the 5’UTR that cause a *lin-28(0)*-like phenotype (Table 1, line 10). These observations suggests that LIN-28 activity post-transcriptionally inhibits *lin-46* expression through direct binding.

**Table 4.**
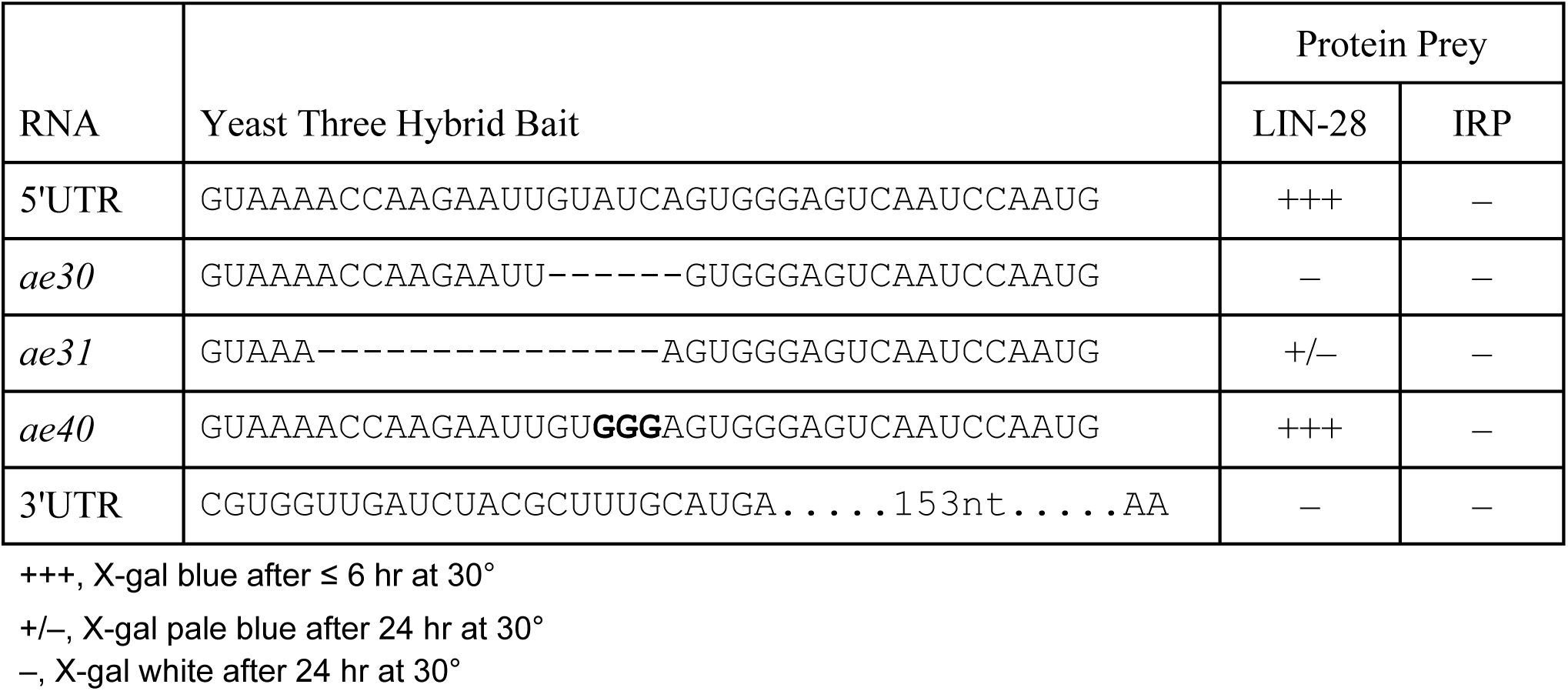
Yeast three-hybrid binding of LIN-28 to *lin-46* 5’UTR.

### LIN-46 binds two of HBL-1’s zinc fingers and inhibits HBL-1 activity in vivo

*lin-46* encodes an unusual protein whose distant relatives are involved in protein-protein interactions, although its mechanism of action in *C. elegans* isn’t known (Pepper et al. 2004). Ilbay and Ambros showed that *lin-46* appears to relocalize the transcription factor HBL-1 from the nucleus to the cytoplasm of seam cells (Ilbay and Ambros 2019). To begin to understand how the LIN-46 protein works at the molecular level, we performed a yeast two-hybrid screen using a library of *C. elegans* cDNAs and identified HBL-1 as a directly interacting protein (<u>Fig. S3</u>). We validated the interaction in vitro by GST pull down (<u>Fig. S3</u>, panel B). Although we cannot rule out the existence of other interacting partners of LIN-46, HBL-1 was the only protein we identified in the screen.

The only other protein we could find that LIN-46 interacts with is itself <u>Fig. S3</u>, panel A). Structural homology to MoeA of *E. coli* suggests that LIN-46 self-associates and that LIN-46 likely interacts with HBL-1 as a dimer (Pepper et al. 2004).

HBL-1 is a Hunchback-like Ikaros family member with nine zinc fingers (Fay et al. 1999). In addition to four zinc fingers involved in DNA binding, this family is characterized by two C-terminal zinc fingers involved in homodimerization and heterodimerization with other family members (McCarty et al. 2003). In yeast two-hybrid assay experiments, LIN-46 interacted exclusively with segments of the protein containing either zinc fingers number 5 or number 9 (<u>Fig. S3</u>, panel C). Zinc finger 5 of HBL-1 is highly conserved among its relatives and is likely involved in DNA binding based on its homology to other family members. Zinc finger 9 is variable in sequence among relatives and is the C-terminal most zinc finger, one of the two ostensibly involved in dimer formation (<u>Fig. S3</u>, panel C). Interestingly, HBL-1 is unique in this family in that it does not homodimerize via its C-terminal zinc fingers (Giesecke et al. 2006). Notably, the allele *hbl-1(ae23)*, which lacks the C-terminal zinc fingers, has no discernable phenotype on its own (Table 1, line 4).

We wondered whether HBL-1 could interact with other Ikaros family members of *C. elegans* under the hypothesis that LIN-46 might disrupt a heterodimerization. We used another yeast-based interaction method that avoids false positives caused by the use of transcription factors as bait (Moerdyk-Schauwecker et al. 2011). We found that whereas HBL-1 could bind to LIN-46 in this assay, we detected no interaction with itself or other *C. elegans* Ikaros family members: SDZ-12, EHN-3, ZTF-16, C46E10.8, and F26F4.8 (data not shown). ZTF-16 is one of these that has been shown to have a role later in the heterochronic pathway than HBL-1 (Hansen et al. 2022). We also screened a library of *C. elegans* sequences for proteins other than LIN-46 that could interact with HBL-1 and found none (data not shown).

Nevertheless, to determine whether LIN-46 is capable of inhibiting HBL-1 activity in vivo, we took advantage of a situation in which *hbl-1* is inappropriately expressed. In some *C. elegans* motor neurons, *hbl-1* is transcriptionally repressed by the transcription factor UNC-55 (Thompson-Peer et al. 2012). When *unc-55* is mutant, HBL-1 is ectopically expressed which leads to inappropriate re-wiring of the synapses of these motor neurons, resulting in the characteristic “uncoordinated” phenotype of *unc-55* mutants. If LIN-46 is an inhibitor of HBL-1 activity in vivo, ectopic expression of LIN-46 in neurons of *unc-55* mutants should suppress the ectopic rewiring and alleviate the uncoordinated phenotype.

We previously reported that a *lin-46::GFP* translational fusion is visible only periodically in the lateral and ventral epidermis, with no discernible fluorescence in the surrounding hypodermal syncytium or other sites (Pepper et al. 2004). We were unsuccessful when attempting to express this fusion protein constitutively or more broadly in hypodermal cells using heterologous promoters (e.g., *col-10*), suggesting that LIN-46 accumulation is restricted temporally and spatially by a post-translational mechanism (<u>Fig S4</u>, panel A). However, we were able to express LIN-46:GFP constitutively in cells of the nervous system using the *rgef-1* promoter (<u>Fig. S4</u>, panel B)(Chen et al. 2011).

*unc-55* mutants have a characteristic coiling behavior when moving backward, and loss-of-function mutations in *hbl-1* partially suppress this phenotype (Thompson-Peer et al. 2012). We found that ectopic expression of LIN-46 in neurons suppressed the *unc-55* mutant’s coiling to the same degree as a *hbl-1* mutation, whereas expressing a paralog of LIN-46 (MOC-1) had no effect (<u>Fig. S5</u>, panel A). Coiling is due to a defect in synapse formation which can be visualized on both the dorsal and ventral sides using reporters. By counting the number of synapses, we determined that ectopic LIN-46 also alters the number of synapses in a *unc-55* mutant to the same degree as loss of *hbl-1* (<u>Fig. S5</u>, panel B).

Finally, to test whether zinc fingers of HBL-1 could bind to and inhibit LIN-46 in vivo, we expressed a C-terminal domain of HBL-1 with LIN-46-binding activity (labeled with an asterisk in <u>Fig. S3</u>), in developing larave using the *col-10* promoter. We found that ectopically expressed HBL-1 C-terminal domain caused a *lin-46(0)*-like phenotype, producing gaps in adult later alae <u>Table S1</u>. The notable difference was that whereas *lin-46(0)* is cold-sensitive, expression of the HBL-1 C-terminal domain was not (<u>Table</u> <u>S1</u>). Our results therefore suggest that LIN-46 directly inhibits HBL-1 activity by binding two of its zinc fingers, 5 and 9, possibly interfering with DNA binding.

### lin-28 acts only in part through lin-46 to have its effect on gonad development

Because the sterility of the *lin-28(0); hbl-1(lf)* double mutant was unlike either mutant alone, it is possible that loss of *lin-28* further reduced the activity of a *hbl-1* hypomorphic allele. Previous work from the lab suggested that *lin-28* might act via *hbl- 1*’s 3’UTR (Vadla et al. 2012). However, because the results presented above suggest *lin-28* regulates *hbl-1* via *lin-46*, we characterized *lin-46*’s role with respect to the *lin-28; hbl-1* gonad phenotype by genetic analysis. Specifically, we sought to determine whether *lin-28* acts entirely or in part through *lin-46* to regulate *hbl-1*.

Mutations in *lin-28* and *lin-46* cause opposite heterochronic phenotypes and fully suppress each other’s timing defect when combined (Pepper et al. 2004). There is a significant difference in the penetrance and expressivity of their phenotypes, indicating that the genes do not act in a simple linear pathway (Pepper et al., 2005). Additionally, alleles of *lin-46* are not able to suppress *hbl-1* hypomorphic allele phenotypes which resemble those of *lin-28* null mutants (Pepper et al. 2004; Abbott et al. 2005). To test whether *lin-46* null would suppress the sterility phenotype of the *lin-28(0); hbl-1(lf)* double mutant, we generated a *lin-28(0); lin-46(0); hbl-1(lf)* strain. We found the sterility phenotype was partially rescued in this strain (Table 1, line 8). Suppression occurred with both weak and strong *hbl-1* alleles, although fertility was not fully restored for either (Table 1, lines 8 and 9). The brood sizes of fertile triple mutants were similar to those seen in *lin-28* and *hbl-1* single mutants.

The *lin-46(ae30)* allele, which has a 6-bp deletion in its 5’UTR that prevents LIN- 28 binding (Table 4) produces a precocious phenotype similar to a *lin-28* null allele (Table 1, line 10). This allele resembles those identified by Ilbay and Ambros (Ilbay et al. 2021). If *lin-28* supports *hbl-1* expression or activity via *lin-46,* then *lin-46(ae30)* when combined with a *hbl-1* hypomorphic allele should cause sterility. *lin-46(ae30); hbl- 1(ve18)* did display slightly lower fertility in comparison to *hbl-1(ve18)* alone, but not as low as the *lin-28(0); hbl-1(ve18)* animals (Table 1, compare lines 3, 5 and 11).

Furthermore, we saw no reduction in fertility when *lin-46(ae30)* was combined with the weaker *hbl-1(ae23)* allele, unlike the *lin-28(0); hbl-1(ae23)* double mutant (Table 1, compare lines 6 and 12). This suggests that a significant fraction of *lin-28*’s effect on *hbl-1* occurs independently of *lin-46*.

### HBL-1 activity is necessary for normal gonad development

Because a null allele of *hbl-1* is embryonic lethal, it was difficult to directly address whether *lin-28* and *hbl-1* act in a linear or a branched pathway to affect gonad development. To resolve this issue, we analyzed gonad development after supplying HBL-1 activity during embryogenesis and removing it during larval development. To control when HBL-1 is active, we utilized the auxin-inducible degron (AID) system where the addition of the plant hormone auxin to nematode growth media causes in vivo degradation of proteins containing the auxin-binding degron (Zhang et al. 2015; Hills-Muckey et al. 2021 Oct 12).

We generated a strain that contained the endogenous *hbl-1* locus tagged at its C-terminus with the degron sequence (*hbl-1::AID*). Without auxin, these animals showed a wildtype phenotype, whereas, when grown on auxin-containing media, they displayed a phenotype like that of the *hbl-1(ve18)* allele where HBL-1 activity is reduced (Table 1, lines 13 and 14). To maximally reduce *hbl-1* activity during larval development, we grew newly hatched *hbl-1::AID* animals on media containing auxin and *E. coli* expressing *hbl-1* dsRNA (*hbl-1* RNAi). Eggs from this strain were transferred onto auxin+RNAi plates at the threefold stage of development and grown to adulthood.

Under these conditions, we observed that 50% of the *hbl-1::AID* animals displayed a sterility phenotype with fertile animals producing a reduced number of offspring (Table 1, line 16). Many sterile animals of adult age were found to display abnormal gonad morphology, like that of the *lin-28(0); hbl-1(lf)* mutants, with no fertilized embryos present. Thus, reduction of *hbl-1* activity alone during larval development is sufficient to affect gonad morphogenesis. Removal of *lin-28* activity from *hbl-1::AID* animals reared on *hbl-1* RNAi and auxin increased the severity of the sterility defect (Table 1, line 17). Nevertheless, these observations support a model whereby *lin-28* regulates the expression and/or activity of *hbl-1*, and *hbl-1* is the proximal gene responsible for normal gonad development.

As demonstrated above, *lin-46* is able to suppress the sterility phenotype of the *lin-28(0); hbl-1(lf)* doubles. We sought to determine whether *lin-46* would still be able to suppress the sterility phenotype of the *lin-28(0); hbl-1::AID* double mutant. The removal of *lin-46* from the *lin-28(0); hbl-1::AID* resulted in a low level of rescue of the sterility phenotype. Fertility was restored to 27% of *lin-28(0); lin-46(0); hbl-1::AID* triple mutants (Table 1, line 18). Additionally, the progeny number in these triple mutants was significantly lower than that seen in *lin-28(0)* and *lin-46(0)* single mutants.

By using the alternative AID system, we were able to see that *lin-46* exerts only a modest rescue of the sterility phenotype. Moreover, the magnitude of rescue by *lin-46* is significantly lower in the *lin-28(0); hbl-1::AID* double mutant than in the *lin-28(0); hbl-1(lf)* double mutants, further indicating *lin-46* is only able to rescue the sterility phenotype if there is HBL-1 protein present. These results support the idea that *lin-28* acts only partially through *lin-46* to exert positive regulation of *hbl-1*. Evidence also supports the model that *hbl-1* is the most proximal acting factor in the heterochronic pathway for the control of somatic gonad development.

## Discussion

Here we report a gonad morphogenesis phenotype the extent of which had not been reported for heterochronic mutants and which is not connected in an obvious way with developmental timing. Furthermore, the genetic relationships among the genes with respect to this phenotype, along with new evidence of their direct interactions, shed some light on so far unresolved molecular relationships among these regulators.

We found that when combining loss-of-function alleles of *lin-28* and *hbl-1* that on their own cause heterochronic phenotypes in the hypodermis, there occurred an unexpected level of sterility due to abnormal gonad growth and development. In addition to abnormal DTC migration seen in some other developmental timing mutants, developmental failure of the hermaphrodite spermatheca is severe (Fig. 1). This phenotype is not merely additive: Whereas single mutants of *lin-28* and *hbl-1* occasionally had one arm of the gonad display abnormal migration, in more that half of the double mutants both arms migrated abnormally, and it was uncommon to see an animal with normal gonad migration (Table 2). Furthermore, neither of the single mutants ever lacked a spermatheca entirely, but again more than half of the double mutants completely lacked one or both structures (Table 3).

The heterochronic genes are characterized by temporal cell fate transformations, substituting one stage-specific fate for another. It is difficult to determine whether cell fate transformations like this occurred in the gonad of *lin-28; hbl-1* double mutants: gonad development appeared normal through the L2 when most the lineages producing gonadal cells complete. The first defect we observed was the aberrant migration of DTCs at the start of the L3. Although *lin-28* and *hbl-1* have not been previously implicated in governing DTC migration, later-acting heterochronic genes that act in parallel with or downstream of both in the heterochronic pathway have been shown to have a significant role in DTC migration in late larval development (Huang et al. 2014; Cecchetelli and Cram 2017). It is believed that these later-acting genes control signaling within the DTCs themselves, so learning where *lin-28* and *hbl-1* are required to affect gonad morphogenesis—whether in the DTCs, the hypodermis, or another site— will be important to understanding this phenomenon.

It is far less clear how gonad morphogenesis, specifically spermathecal and uterine development, are governed by heterochronic gene activity since single gene mutants with strong hypodermal phenotypes show no such defects. Again, whether these genes act within the hypodermis to affect gonad morphogenesis or within the gonad itself must be determined. Still, any theory explaining how the early-acting heterochronic genes impact gonad development of the hermaphrodite must take into account that male gonad development appears normal, despite their high-penetrance impacts on hypodermal development in males.

We initiated these studies to clarify the relative roles of *lin-28*, *lin-46*, and *hbl-1* in the heterochronic pathway. Previous genetic studies indicate that the activities of multiple factors of the heterochronic pathway converge onto *hbl-1*, which appears to be the most proximal acting gene involved in controlling cell fates of the L2 (Abbott et al. 2005; Ilbay and Ambros 2019). Indeed, we provide new evidence for direct interactions between the RNA-binding protein LIN-28 and the 5’UTR of *lin-46* (Table 4), and between LIN-46 and HBL-1 zinc fingers (<u>Fig. S3</u>), also confirming that LIN-46 can inhibit HBL-1 activity in vivo (<u>Fig. S5</u>). Along with prior genetic evidence, these data are consistent with a model whereby *lin-28* acts via negative regulation of *lin-46* to positively regulate *hbl-1* in a linear pathway. However, other lines of evidence indicate that that model does not account for a *lin-46-*independent regulation of *hbl-1* by *lin-28* (Pepper et al. 2004; Vadla et al. 2012). Here we find further indications that *lin-28* does not act solely through *lin-46* to regulate *hbl-1*. Particularly, a mutation in the 5’UTR of *lin-46* that abrogates LIN-28 binding does not phenocopy the *lin-28(0); hbl-1(lf)* sterility in that most of the *lin-46(utr); hbl-1(lf)* animals were fertile with normal gonads (Table 1, compare lines 5 and 11). We conclude, therefore, that *lin-28* acts in a forked fashion to influence *hbl-1* activity, possibly post-transcriptionally (via its 3’UTR) and post-translationally (via LIN-46). Further studies could assess the relative weights of these two branches.

Our results show that *hbl-1* is required during larval development for normal gonad morphogenesis. The work of Ilbay and Ambros and our own unpublished findings show that deletion of microRNA regulatory sites in the 3’UTR of *hbl-1* does not impact gonad development (Ilbay and Ambros 2019). This difference suggests that the mere presence of HBL-1 and not its temporal regulation matters for normal fertility. The threshold for the level of HBL-1 activity required for normal somatic gonad development appears to be lower than the level of activity required for correct developmental timing in the hypodermis as hypomorphic mutants produce a heterochronic effect while gonad development is normal.

*hbl-1*, which encodes an Ikaros-family transcription factor, is known to be pleiotropic in *C. elegans* development. In addition to its larval developmental timing function, *hbl-1* has been shown to have roles in late embryonic development, synaptic remodeling, and dauer formation (Fay et al. 1999; Karp and Ambros 2011; Thompson-Peer et al. 2012). Where these roles differ from *hbl-1*’s effect on the gonad is that we see a role for other heterochronic genes in the later. In other words, we believe that normal gonad development does not merely require *hbl-1* and no other heterochronic pathway members like, say, late embryonic hypodermal development. Rather, gonad development requires multiple members of at least one branch of the heterochronic pathway, suggesting a more extensive role for the developmental timing genes in gonad development than previously known.

Finally, we recently observed that *Caenorhabditis briggsae* orthologs of *C. elegans* heterochronic genes show mutant phenotypes that differ in significant ways from those of *C. elegans* (Ivanova and Moss 2023). Specifically, several of these *C. briggsae* mutants show a gonad disintegration phenotype that closely resembles the gonad phenotype we describe here. The *C. briggsae* genes with apparent roles in gonad development include *Cbr-lin-28*, *Cbr-lin-46*, *Cbr-hbl-1*, and *Cbr-lin-41*. Compared to their *C. elegans* counterparts, these genes show much weaker heterochronic phenotypes in the hypodermis, whereas their roles in gonad development are more apparent, suggesting significant evolutionary drift in their activities since the last common ancestor of *C. elegans and C. briggsae*. Nevertheless, despite this drift, we note that the heterochronic genes of both species have clear roles in both developmental timing and gonad morphogenesis, suggesting an enduring association between these aspects of development.

## Materials and Methods

### Culture conditions

Nematodes were grown under standard conditions at 20°C unless otherwise indicated (Sulston and Horvitz 1977). Strains with multiple allelic mutations were generated by following lesions via PCR and confirmed through DNA sequencing. Most strains contain the wIs78 transgene, a derivative of wIs1, which contains seam cell nuclei (*scm::GFP*) and seam cell junction (*ajm-1::GFP*) markers to identify and quantify lateral hypodermal seam cells (Koh and Rothman 2001). Males for gonad analysis were produced by growing strains on *E. coli* bacteria containing the RNAi plasmid pLT651 (Timmons et al. 2014).

### Strains used

N2 wild type

RG733 *wIs78 (ajm-1::GFP & scm::GFP)*

ME414 *wIs78; hbl-1(ae23) X*

ME419 *wIs78; lin-28(ga54) I*

ME423 *wIs78; lin-28(ga54) I; hbl-1(ae26) X*

ME429 *wIs78; hbl-1(ae26) X*

ME433 *wIs78; lin-28(ga54) I; lin-46(ma174) V; hbl-1(ae23) X*

ME438 *wIs78; lin-46(ae30) V*

ME443 *wIs78; lin-28(ga54) I; hbl-1(ae23)/tmC30[myo-2p::Venus]X*

ME452 *wIs78; lin-28(ga54) I; hbl-1(ve18)/tmC30[myo-2p::Venus]X*

ME455 *wIs78; lin-28(ga54) I; qIs56[lag-2p::GFP + unc-119(+)] V; hbl-1(ae23)/ tmC30[myo-2p::Venus]X*

ME456 *wIs78; lin-28(ga54) I; ezIs2[fkh-6::GFP + unc-119(+)] III; hbl-1(ae23)/ tmC30[myo-2p::Venus]X*

ME481 *wIs78; lin-28(ga54) I; qIs56[lag-2p::GFP + unc-119(+)] V*

ME485 *wIs78; lin-28(ga54) I; ezIs2[fkh-6::GFP + unc-119(+)] III*

ME502 *cshIs140 [rps-28pro::TIR-1(F79G)_P2A mCherry-His-11;*

*Cbunc-119(+)] II; HBL-1-AID (hbl-1 (aeIS8 (12.4), HBL-1::AID)); wIs78 (pDP#MM016B (unc-119) + pJS191 (AJM-1::GFP + pMF1 (SCM::GFP) + F58E10)*

ME509 *wIs78; lin-28(ga54) I; arTi145 [ckb-3p::mCherry::his-58::unc-54 3’UTR] II; hbl- 1(ve18)/tmC30[myo-2p::Venus]X*

ME521 *wIs78; lin-28(ga54) I; cshIs140 [rps-28pro::TIR-1(F79G)_P2A mCherry- His-11; Cbunc-119(+)] II; HBL-1-AID (hbl-1 (aeIS8 (12.4), HBL-1::AID))*

ME522 *wIs78; lin-28(ga54) I; lin-46(ma174) V; cshIs140 [rps-28pro::TIR-1(F79G)_P2A mCherry-His-11; Cbunc-119(+)] II; HBL-1-AID (hbl-1 (aeIs8 (12.4), HBL-1::AID))* ME524 *hbl-1(ve18)X; ezIs2[fkh-6::GFP + unc-119(+)] III*

ME525 *hbl-1(ve18) X; qIs56[lag-2p::GFP + unc-119(+)] V*

ME542 *wIs78; lin-46(ae30) V; hbl-1(ae23) X*

ME543 *wIs78; lin-46(ae30) V; hbl-1(ve18) X*

ME551 *wIs78; lin-28(ga54) I; lin-46(ma174) V; hbl-1(ve18) X*

ME568 *unc-55(e1170); nuIs279 aeEx49 (Punc-25:lin-46, ttx-3::GFP)* ME569 *unc-55(e1170); nuIs279 aeEx50 (Punc-25:moc-1, ttx-3::GFP)* KP5348 *nuIs279 [Punc-25:unc-57:GFP + Punc-25:mCherry:rab-3]* KP5363 *unc-55(e1170); nuIs279*

KP6119 *unc-55(e1170); hbl-1(mg285); nuIs279*

### Microscopy

Nomarski DIC and fluorescence microscopy were used for quantitative and qualitative assessments of phenotypes. Developmental stage was assessed by time from hatching, extent of gonad development, and in some cases counting molts. Images were acquired with AxioCam with AxioVision with Plan-NEOFLUAR 63x and alpha Plan-FLUAR 100x objectives on a Zeiss Axioplan2 microscope.

### RNA interference

Bacterial-mediated RNA-interference was performed as previously described (Timmons et al. 2014). The RNAi vectors used contain a 3.5kb region of *hbl-1* genomic sequence in the T444T backbone (Sturm et al. 2018). Bacteria were induced in culture and seeded on nematode growth media (NGM) plates containing 1mM ITPG (GoldBio) and 50μg/ml ampicillin (VWR Life Science) and 12.5 μg/ml tetracycline (FisherBiotech).

### Auxin-inducible Degron

The AID-tagged *hbl-1* allele was constructed using the CRISPR/Cas9 as described using the pDD162 vector and *dpy-5* as a co-CRISPR marker (Dickinson and Goldstein 2016; El Mouridi et al. 2017). The gRNA was synthesized using InvitrogenTM, MEGAshortscriptTM T7 Transcription Kit and the repair template was generated primers:

5’ ATCAGGCTTTCAATGAGCTAAGCTTTGCTCTCCACATGTAtCAgGCgcGtCAtC

Agatgcctaaagatccagccaaacc

5’ TTAACGAGGACGTCCTCATTATTGGTGTCTGGCTTGGTACTTActtcacgaacgccg ccgc

Repair templates were mixed and heated to 96°C for 5 minutes and then placed on ice (Dokshin et al. 2018). The sequence targeted the 3’ end of the *hbl-1* protein coding region. The guide RNA sequence used targeted the top strand: 5’ CATGTACCAAGCCAGACACC. 5-Ph-IAA was resuspended in DMSO to make 10mM stocks (Hills-Muckey et al. 2021). Animals were grown on two variations of NGM plates. Plates treated with 1μM 5-Ph-IAA (Sigma-Aldrich), or plates treated with: 1μM 5-Ph-IAA, 1mM ITPG (GoldBio), 50μg/ml ampicillin (VWR Life Science), and 12.5 μg/ml tetracycline (FisherBiotech).

### Yeast two- and three-hybrid assays

Yeast two-hybrid assays were performed as described previously (Golemis et al. 2008). The *lin-46* open reading frame was fused to the DNA-binding domain in pMW103. Portions of the *hbl-1* open reading frame were fused to the activation domain in pJG4-5. These plasmids were co-transformed with the *lacZ* reporter. X-gal overlays were assessed after 6 hours and overnight.

Yeast three-hybrid assays were performed using the YBZ-1 strain as described previously (Hook et al. 2005). The *lin-28* open reading frame was fused to the activation domain sequence in pACT2, and experimental RNAs were fused to the MS2 stem loop sequence in pIIIA/MS2-2. X-gal overlays were assessed after 6 hours and overnight.

## Data availability

Strains and plasmids are available upon request. Figures and supplemental files available at figshare; https://doi.org/xxx.

## Acknowledgements

The authors thank Disha Patel for generating the *lin-46* 5’UTR mutants. We thank WormBase for genetic sequences and WormAtlas for reference information on nematode biology and development. Some strains were provided by the CGC, which is funded by NIH Office of Research Infrastructure Programs (P40 OD010440).

---

**Fig. S1.**
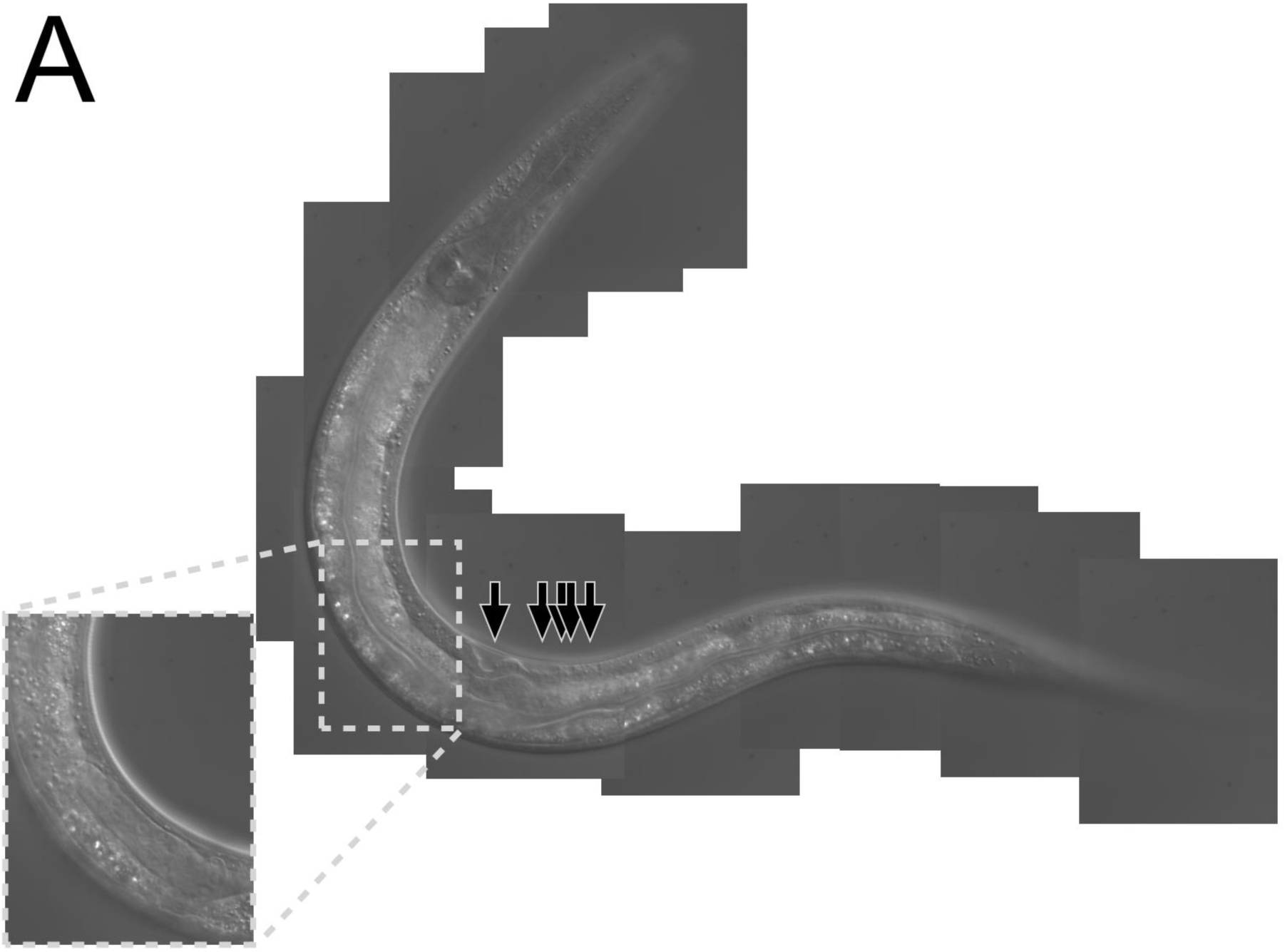
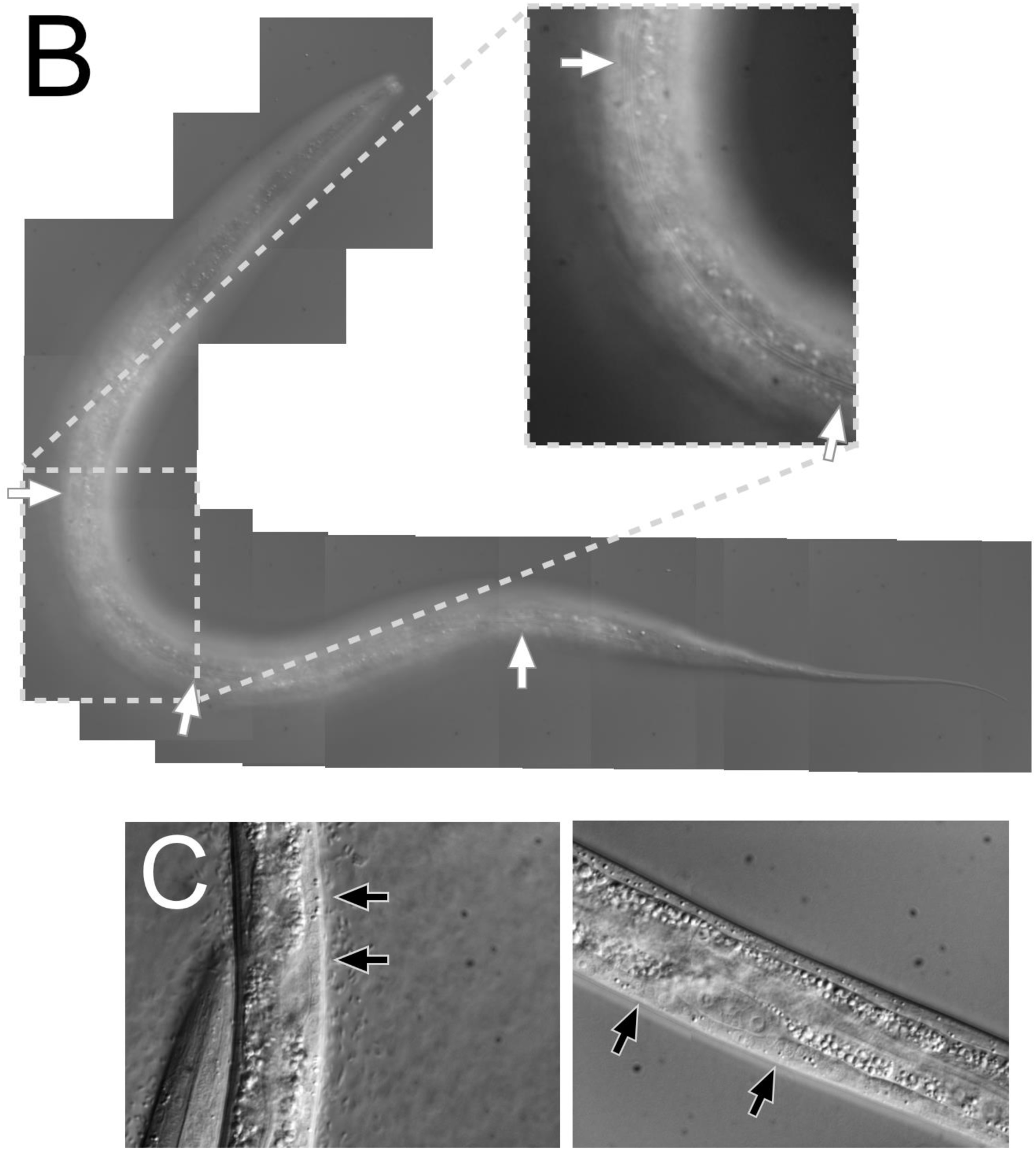
Phenotypes displayed by *lin-28; hbl-1* larvae. **A.** *lin-28; hbl-1 L3* larvae displaying advanced vulva morphogenesis and abnormal multivulva (Muv) morphology (black arrows). **Inset Box** showing stage in larval development based on gonad arm position. **B.** *lin-28; hbl-1* L3 larvae displaying full adult-specific lateral alae (white arrows). **Inset Box** showing magnified view of alae. **C.** Individual lin-28; hbl-1 larvae at L2 molt displaying precocious VPC divisions (black arrows).

**Fig. S2.**
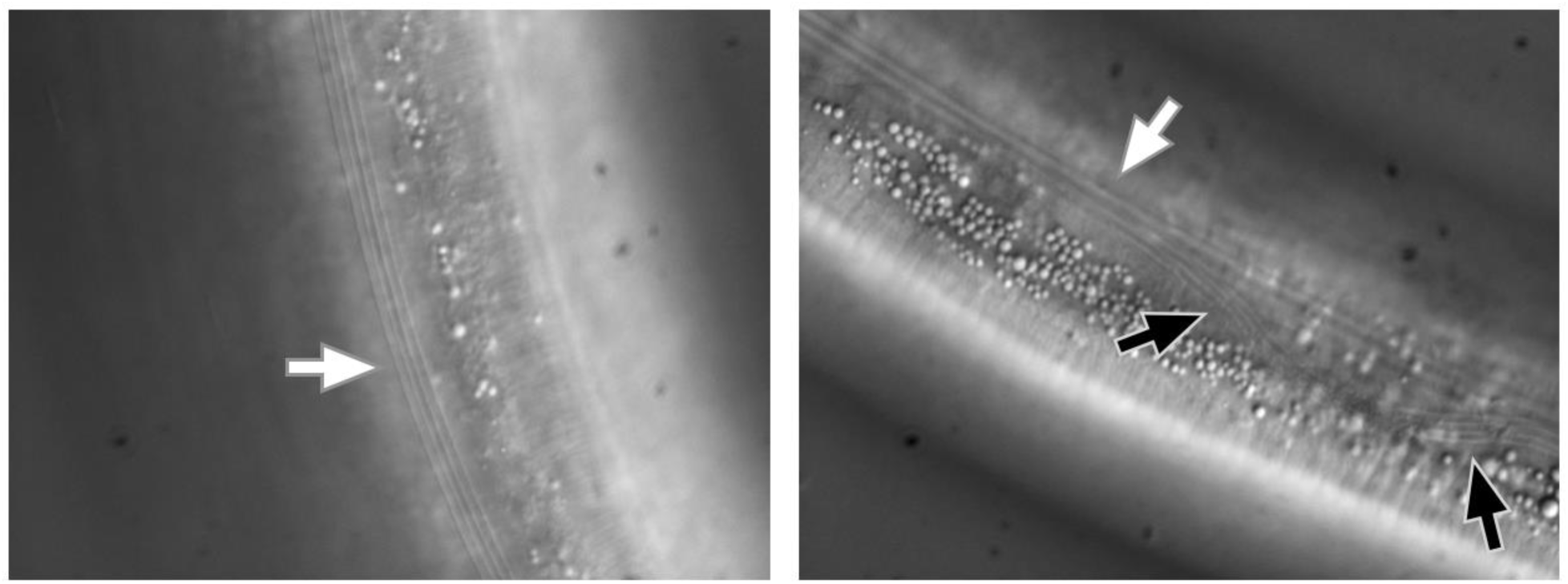
Molting defects displayed by *lin-28; hbl-1* mutant. **A.** Wild type *C. elegans* adult displaying alae on single cuticle (white arrow). **B.** A similarly-aged *lin-28; hbl-1* double mutant displaying alae ridges from older, unshed cuticle (white arrow) as well as a newer cuticle that can be seen underneath (black arrows).

**Fig. S3.**
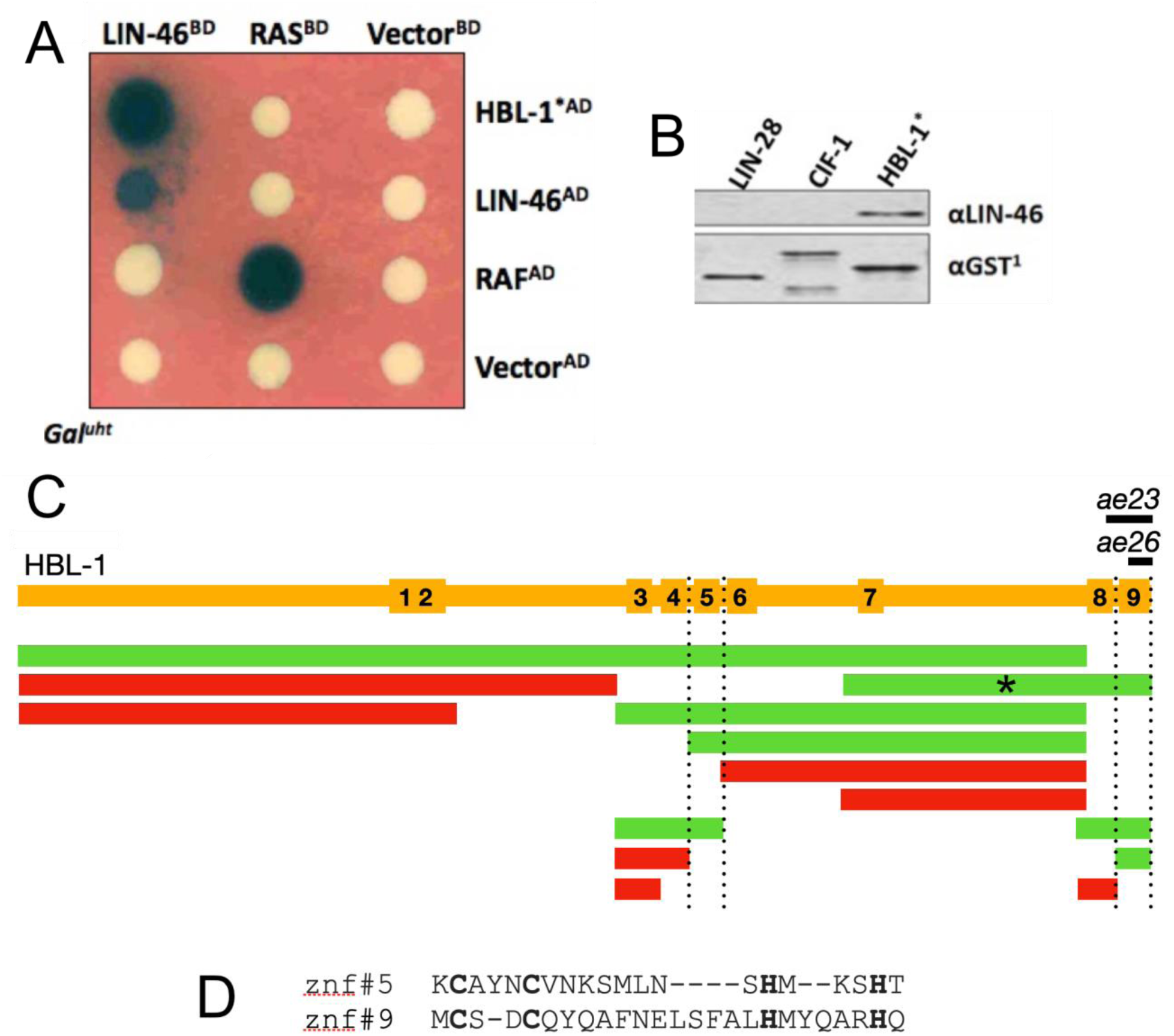
Interaction of HBL-1 with LIN-46. A, Yeast two-hybrid tests of binding among LIN-46, HBL-1, and control proteins (RAS and RAF). HBL-1* is a C-terminal fragment of HBL-1 indicated by the asterisk in panel C. Dark blue colonies are indicative of an interaction, white means no interaction. BD, DNA binding domain. AD, activation domain. **B,** Immunoblots of a GST pull-down experiment where proteins labeled across the top were fused with GST and mixed with unfused LIN-46. Bottom panel: All three GST fusions could be detected in the pull-down by anti-GST antisera. Top panel: Only the HBL-1* fragment pulled down LIN-46. **C,** Summary of yeast two-hybrid interactions between LIN-46 and portions of HBL-1. A schematic of the HBL-1 protein sequence showing the locations of 9 zinc fingers and indicating the sequences missing in alleles *ae23* and *ae26* (black bars). Green: HBL-1 portions that show a strong positive interaction with LIN-46 in a yeast two-hybrid assay. Red: Portions that show no interaction with LIN-46. **D,** The sequences of zinc fingers 5 and 9 aligned by their cysteine and histidine residues showing their dissimilarity.

**Fig. S4.**
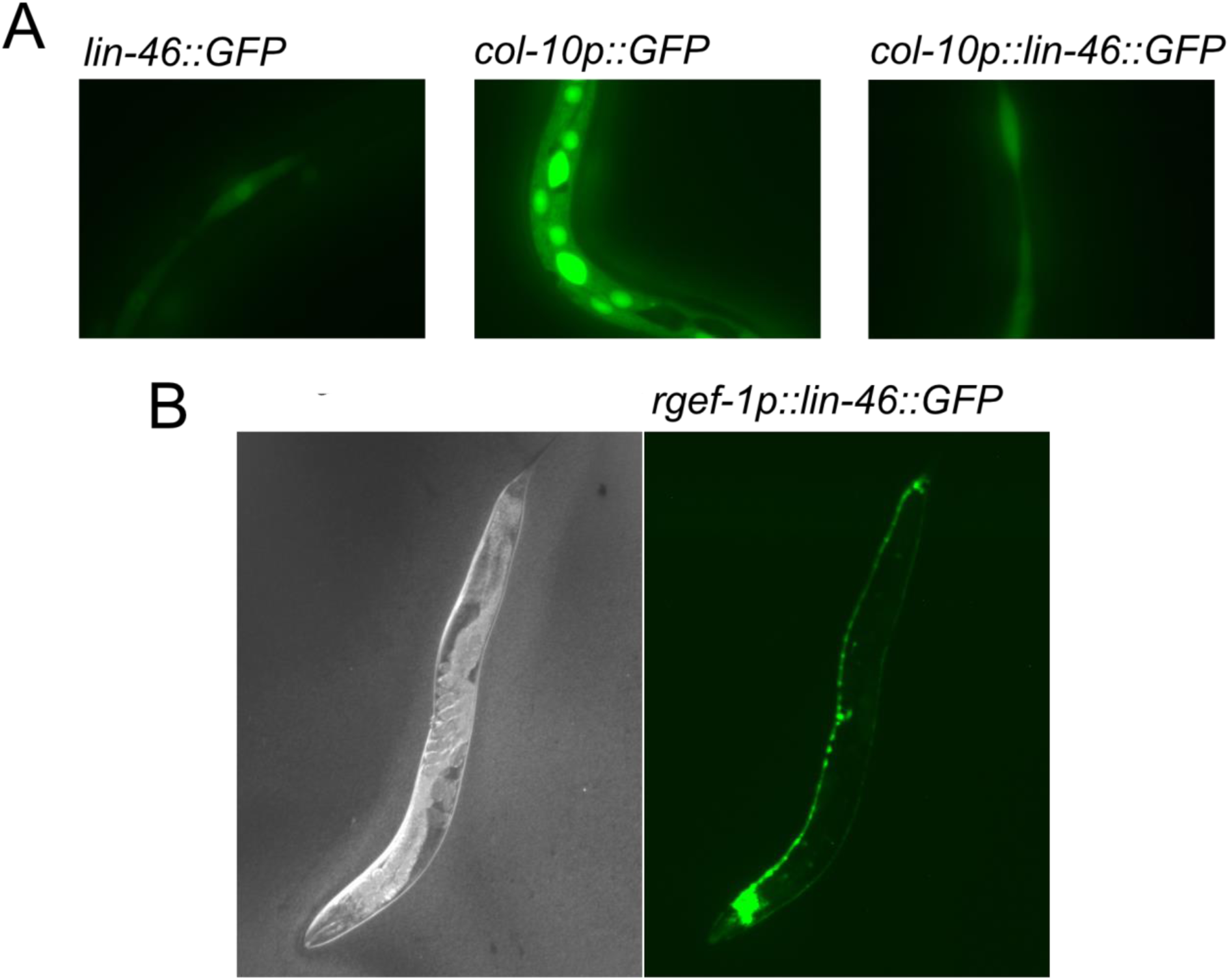
Ectopic expression of LIN-46. A, Fluorescence micrographs of late-stage larvae expressing the indicated transgenes. For both *lin-46::GFP* and *col-10p::lin-46::GFP*, fluorescence is seen dimly in seam cells at the time of the molts, whereas *col-10p::GFP* is bright continuously throughout the hypodermis. **B,** A DIC image (left) and a fluorescence image (right) of an adult expressing *rgef-1::lin-46::GFP*. Expression is continuous throughout the nervous system.

**Fig. S5.**
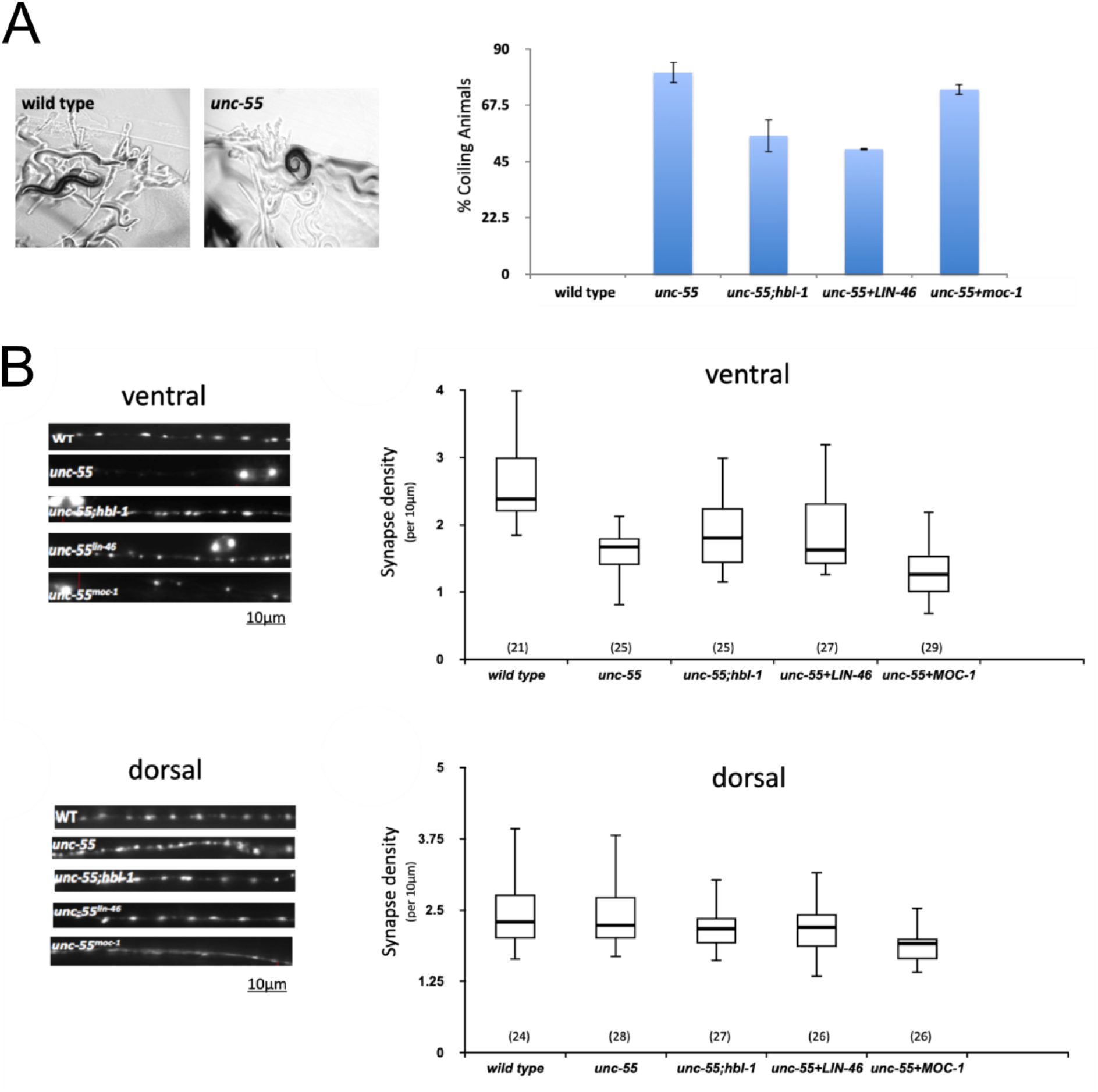
LIN-46 inhibits HBL-1 activity in vivo. A, Left, A comparison of the movement of wildtype and *unc-55* mutant animals. Wild type backs up in a sinusoidal motion, whereas *unc-55* mutants move backward with difficulty and frequently coil. A *hbl-1* mutation decreases the coiling of *unc-55* mutants (Thompson-Peer et al. 2012). Right, Percentages of animals of different genotypes that display coiling behavior. LIN-46 and MOC-1 (a LIN-46 paralog) were ectopically expressed in *unc-55* mutants. **B,** *unc-55* mutants have fewer ventral synapses but the normal amount of dorsal synapses. A *hbl-1* mutation increases the number of ventral synapses of *unc-55* mutants (Thompson-Peer et al. 2012). Synapses are visualized using the endophilin marker *unc-57:GFP*. Left, representative images of ventral and dorsal synapses in strains of different genotypes. Right, quantitation of synapse density per 10 microns of strains of different genotypes.

**Table S1.** *lin-46(0)-*like phenotype caused by ectopic HBL-1 C-terminal fragment.

|  | Genotype and growth temperature | % with gaps in adult alae (*n*)<sup>a,b</sup> |
| --- | --- | --- |
|  | wild type |  |
| 1 | 15° | 0 (20) |
| 2 | 20° | 0 (20) |
| 3 | 25° | 0 (22) |
|  | *lin-46(0)* |  |
| 4 | 15° | 100 (20) |
| 5 | 20° | 29 (24) |
| 6 | 25° | 4 (24) |
|  | *col-10p::hbl-1C-term.* |  |
| 7 | 15° | 5 (21) |
| 8 | 20° | 45 (20) |
| 9 | 25° | 23 (22) |

<sup>a</sup> Alae gaps observed in young adult animals covered one to several seam cells.
<sup>b</sup> Number of animals (*n*) scored for each genotype is indicated in parenthesis; one side scored per animal.

## References

1. Abbott AL, Alvarez-Saavedra E, Miska EA, Lau NC, Bartel DP, Horvitz HR, Ambros V. 2005. The let-7 MicroRNA family members mir-48, mir-84, and mir-241 function together to regulate developmental timing in Caenorhabditis elegans. Dev Cell. 9(3):403–414. doi:10.1016/j.devcel.2005.07.009.

2. Abrahante JE, Daul AL, Li M, Volk ML, Tennessen JM, Miller EA, Rougvie AE. 2003. The Caenorhabditis elegans hunchback-like gene lin-57/hbl-1 controls developmental time and is regulated by microRNAs. Dev Cell. 4(5):625–637. doi:10.1016/s1534-5807(03)00127-8.

3. Ambros V, Horvitz HR. 1984. Heterochronic mutants of the nematode Caenorhabditis elegans. Science. 226(4673):409–416. doi:10.1126/science.6494891.

4. Antebi A, Culotti JG, Hedgecock EM. 1998. daf-12 regulates developmental age and the dauer alternative in Caenorhabditis elegans. Dev Camb Engl. 125(7):1191–1205.

5. Blelloch R, Kimble J. 1999. Control of organ shape by a secreted metalloprotease in the nematode Caenorhabditis elegans. Nature. 399(6736):586–590. doi:10.1038/21196.

6. Cecchetelli AD, Cram EJ. 2017. Regulating distal tip cell migration in space and time. Mech Dev. 148:11–17. doi:10.1016/j.mod.2017.04.003.

7. Chang W, Tilmann C, Thoemke K, Markussen F-H, Mathies LD, Kimble J, Zarkower D. 2004. A forkhead protein controls sexual identity of the C. elegans male somatic gonad. Dev Camb Engl. 131(6):1425–1436. doi:10.1242/dev.01012.

8. Chen L, Fu Y, Ren M, Xiao B, Rubin CS. 2011. A RasGRP, C. elegans RGEF-1b, couples external stimuli to behavior by activating LET-60 (Ras) in sensory neurons. Neuron. 70(1):51–65. doi:10.1016/j.neuron.2011.02.039.

9. Choi S, Ambros V. 2019. The C. elegans heterochronic gene lin-28 coordinates the timing of hypodermal and somatic gonadal programs for hermaphrodite reproductive system morphogenesis. Dev Camb Engl. 146(5). doi:10.1242/dev.164293.

10. Dickinson DJ, Goldstein B. 2016. CRISPR-Based Methods for Caenorhabditis elegans Genome Engineering. Genetics. 202(3):885–901. doi:10.1534/genetics.115.182162.

11. Dokshin GA, Ghanta KS, Piscopo KM, Mello CC. 2018. Robust genome editing with short single-stranded and long, partially single-stranded DNA donors in Caenorhabditis elegans. Genetics. 210(3):781–787.

12. El Mouridi S, Lecroisey C, Tardy P, Mercier M, Leclercq-Blondel A, Zariohi N, Boulin T. 2017. Reliable CRISPR/Cas9 Genome Engineering in Caenorhabditis elegans Using a Single Efficient sgRNA and an Easily Recognizable Phenotype. G3 Bethesda Md. 7(5):1429–1437. doi:10.1534/g3.117.040824.

13. Euling S, Ambros V. 1996. Heterochronic genes control cell cycle progress and developmental competence of C. elegans vulva precursor cells. Cell. 84(5):667– 676. doi:10.1016/s0092-8674(00)81045-4.

14. Fay DS, Stanley HM, Han M, Wood WB. 1999. A Caenorhabditis elegans homologue of hunchback is required for late stages of development but not early embryonic patterning. Dev Biol. 205(2):240–253. doi:10.1006/dbio.1998.9096.

15. Giesecke AV, Fang R, Joung JK. 2006. Synthetic protein-protein interaction domains created by shuffling Cys2His2 zinc-fingers. Mol Syst Biol. 2:2006.2011. doi:10.1038/msb4100053.

16. Golemis EA, Serebriiskii I, Finley RLJ, Kolonin MG, Gyuris J, Brent R. 2008. Interaction trap/two-hybrid system to identify interacting proteins. Curr Protoc Mol Biol. Chapter 20:Unit 20.1. doi:10.1002/0471142727.mb2001s82.

17. Gupta BP, Hanna-Rose W, Sternberg PW. 2012 Nov 30. Morphogenesis of the vulva and the vulval-uterine connection. WormBook Online Rev C Elegans Biol.:1–20. doi:10.1895/wormbook.1.152.1.

18. Hammell CM, Karp X, Ambros V. 2009. A feedback circuit involving let-7-family miRNAs and DAF-12 integrates environmental signals and developmental timing in Caenorhabditis elegans. Proc Natl Acad Sci U S A. 106(44):18668–18673. doi:10.1073/pnas.0908131106.

19. Hansen MA, Dahal A, Bernstein TA, Kohtz C, Ali S, Daul AL, Montoye E, Panzade GP, Alessi AF, Flibotte S, et al. 2022 Jan 1. ztf-16 is a novel heterochronic modulator that opposes adult cell fate in dauer and continuous life histories in Caenorhabditis elegans . bioRxiv.:2022.06.20.496913. doi:10.1101/2022.06.20.496913. http://biorxiv.org/content/early/2022/06/22/2022.06.20.496913.abstract.

20. Hills-Muckey K, Martinez MAQ, Stec N, Hebbar S, Saldanha J, Medwig-Kinney TN, Moore FEQ, Ivanova M, Morao A, Ward JD, et al. 2021 Oct 12. An engineered, orthogonal auxin analog/AtTIR1(F79G) pairing improves both specificity and efficacy of the auxin degradation system in Caenorhabditis elegans. Genetics. doi:10.1093/genetics/iyab174.

21. Hook B, Bernstein D, Zhang B, Wickens M. 2005. RNA-protein interactions in the yeast three-hybrid system: affinity, sensitivity, and enhanced library screening. RNA N Y N. 11(2):227–233. doi:10.1261/rna.7202705.

22. Huang T-F, Cho C-Y, Cheng Y-T, Huang J-W, Wu Y-Z, Yeh AY-C, Nishiwaki K, Chang S-C, Wu Y-C. 2014. BLMP-1/Blimp-1 regulates the spatiotemporal cell migration pattern in C. elegans. PLoS Genet. 10(6):e1004428. doi:10.1371/journal.pgen.1004428.

23. Ilbay O, Ambros V. 2019. Regulation of nuclear-cytoplasmic partitioning by the lin-28- lin-46 pathway reinforces microRNA repression of HBL-1 to confer robust cell- fate progression in C. elegans. Dev Camb Engl. 146(21). doi:10.1242/dev.183111.

24. Ilbay O, Nelson C, Ambros V. 2021. C. elegans LIN-28 controls temporal cell fate progression by regulating LIN-46 expression via the 5’ UTR of lin-46 mRNA. Cell Rep. 36(10):109670. doi:10.1016/j.celrep.2021.109670.

25. Ivanova M, Moss EG. 2023. Orthologs of the Caenorhabditis elegans heterochronic genes have divergent functions in Caenorhabditis briggsae. Genetics. 225(4). doi:10.1093/genetics/iyad177.

26. Karp X, Ambros V. 2011. The Developmental Timing Regulator hbl-1 Modulates the Dauer Formation Decision in Caenorhabditis elegans. Genetics. 187(1):345–353. doi:10.1534/genetics.110.123992. [accessed 2024 Nov 9]. 10.1534/genetics.110.123992.

27. Koh K, Rothman JH. 2001. ELT-5 and ELT-6 are required continuously to regulate epidermal seam cell differentiation and cell fusion in C. elegans. Dev Camb Engl. 128(15):2867–2880. doi:10.1242/dev.128.15.2867.

28. Kovacevic I, Cram EJ. 2010. FLN-1/filamin is required for maintenance of actin and exit of fertilized oocytes from the spermatheca in C. elegans. Dev Biol. 347(2):247–257. doi:10.1016/j.ydbio.2010.08.005.

29. Lin S-Y, Johnson SM, Abraham M, Vella MC, Pasquinelli A, Gamberi C, Gottlieb E, Slack FJ. 2003. The C. elegans hunchback homolog, hbl-1, controls temporal patterning and is a probable microRNA target. Dev Cell. 4(5):639–650.

30. McCarty AS, Kleiger G, Eisenberg D, Smale ST. 2003. Selective dimerization of a C2H2 zinc finger subfamily. Mol Cell. 11(2):459–470. doi:10.1016/s1097-2765(03)00043-1.

31. Moerdyk-Schauwecker M, Destephanis D, Hastie E, Grdzelishvili VZ. 2011. Detecting protein-protein interactions in vesicular stomatitis virus using a cytoplasmic yeast two hybrid system. J Virol Methods. 173(2):203–212. doi:10.1016/j.jviromet.2011.02.006.

32. Moss EG. 2007. Heterochronic genes and the nature of developmental time. Curr Biol CB. 17(11):R425–434. doi:10.1016/j.cub.2007.03.043.

33. Moss EG, Lee RC, Ambros V. 1997. The cold shock domain protein LIN-28 controls developmental timing in C. elegans and is regulated by the lin-4 RNA. Cell. 88(5):637–646. doi:10.1016/s0092-8674(00)81906-6.

34. Pepper AS-R, McCane JE, Kemper K, Yeung DA, Lee RC, Ambros V, Moss EG. 2004. The C. elegans heterochronic gene lin-46 affects developmental timing at two larval stages and encodes a relative of the scaffolding protein gephyrin. Dev Camb Engl. 131(9):2049–2059. doi:10.1242/dev.01098.

35. Rougvie AE, Moss EG. 2013. Developmental transitions in C. elegans larval stages. Curr Top Dev Biol. 105:153–180. doi:10.1016/B978-0-12-396968-2.00006-3.

36. Sturm Á, Saskoi É, Tibor K, Weinhardt N, Vellai T. 2018. Highly efficient RNAi and Cas9-based auto-cloning systems for C. elegans research. Nucleic Acids Res. 46(17):e105. doi:10.1093/nar/gky516.

37. Sulston JE, Horvitz HR. 1977. Post-embryonic cell lineages of the nematode, Caenorhabditis elegans. Dev Biol. 56(1):110–156. doi:10.1016/0012-1606(77)90158-0.

38. Tenen CC, Greenwald I. 2019. Cell Non-autonomous Function of daf-18/PTEN in the Somatic Gonad Coordinates Somatic Gonad and Germline Development in C. elegans Dauer Larvae. Curr Biol CB. 29(6):1064–1072.e8. doi:10.1016/j.cub.2019.01.076.

39. Tennessen JM, Gardner HF, Volk ML, Rougvie AE. 2006. Novel heterochronic functions of the Caenorhabditis elegans period-related protein LIN-42. Dev Biol. 289(1):30–43. doi:10.1016/j.ydbio.2005.09.044.

40. Thompson-Peer KL, Bai J, Hu Z, Kaplan JM. 2012. HBL-1 patterns synaptic remodeling in C. elegans. Neuron. 73(3):453–465. doi:10.1016/j.neuron.2011.11.025.

41. Timmons L, Luna H, Martinez J, Moore Z, Nagarajan V, Kemege JM, Asad N. 2014. Systematic comparison of bacterial feeding strains for increased yield of Caenorhabditis elegans males by RNA interference-induced non-disjunction. FEBS Lett. 588(18):3347–3351. doi:10.1016/j.febslet.2014.07.023.

42. Vadla B, Kemper K, Alaimo J, Heine C, Moss EG. 2012. lin-28 controls the succession of cell fate choices via two distinct activities. PLoS Genet. 8(3):e1002588. doi:10.1371/journal.pgen.1002588.

43. Zhang L, Ward JD, Cheng Z, Dernburg AF. 2015. The auxin-inducible degradation (AID) system enables versatile conditional protein depletion in C. elegans. Development. 142(24):4374–4384.

